# RASLseqTools: open-source methods for designing and analyzing <u>R</u>NA-mediated oligonucleotide <u>A</u>nnealing, <u>S</u>election, and, <u>L</u>igation <u>seq</u>uencing (RASL-seq) experiments

**DOI:** 10.1101/036061

**Authors:** Erick R. Scott, H. Benjamin Larman, Ali Torkamani, Nicholas J. Schork, Nathan Wineinger, Max Nanis, Ryan Thompson, Reza B. Beheshti Zavareh, Luke L. Lairson, Peter G. Schultz, Andrew I. Su

**Author notes:** To whom correspondence should be addressed.; Correspondence may also be addressed to Erick R. Scott. Present Address: [Erick R. Scott], Icahn Institute for Genomics & Multiscale Biology, Mount Sinai School of Medicine, New York, NY, 10029; [H. Benjamin Larman], Department of Pathology, Johns Hopkins School of Medicine, Baltimore, MD, 21205, USA.

## Abstract

**R**NA-mediated oligonucleotide **A**nnealing, **S**election, and **L**igation (RASL-seq) is a method to measure the expression of hundreds of genes in thousands of samples for a fraction of the cost of competing methods. However, enzymatic inefficiencies of the original protocol and the lack of open source software to design and analyze RASL-seq experiments have limited its widespread adoption. We recently reported an Rnl2-based RASL-seq protocol (RRASL-seq) that offers improved ligation efficiency and a probe decoy strategy to optimize sequencing usage. Here, we describe an open source software package, RASLseqTools, that provides computational methods to design and analyze RASL-seq experiments. Furthermore, using data from a large RRASL-seq experiment, we demonstrate how normalization methods can be used for characterizing and correcting experimental, sequencing, and alignment error. We provide evidence that the three principal predictors of RRASL-seq reproducibility are barcode/probe sequence dissimilarity, sequencing read depth, and normalization strategy. Using dozens of technical and biological replicates across multiple 384-well plates, we find simple normalization strategies yield similar results to more statistically complex methods.

## INTRODUCTION

Gene expression signature profiling has found widespread adoption in a number of biological and medical applications^1,2^. However current methods to assess gene expression signatures are either too expensive or too cumbersome to be applied to large-scale patient sampling or high throughput drug discovery. **R**nl2-based **R**NA **A**nnealing, **S**election, and **L**igation-**seq**uencing (RRASL-seq, Figure 1) provides a scalable and cost-efficient method to profile the expression of more than one-hundred transcripts for less than one dollar per sample, thereby facilitating large scale drug discovery screens^3^.

**Figure 1.**
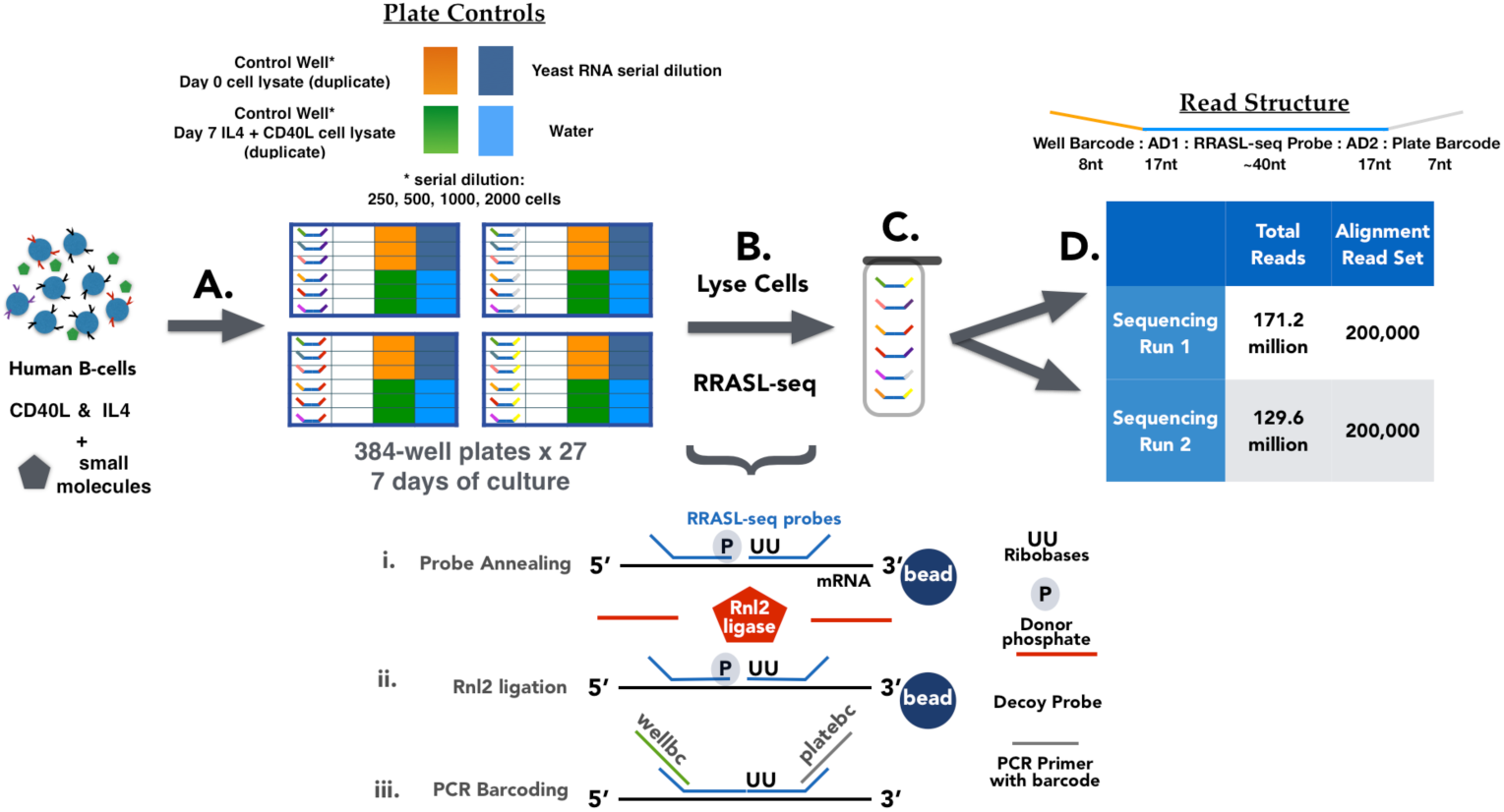
Human IgE B Cell Screen: 1) Human B cells were cultured in IgE polarizing conditions (IL4, CD40L) for 7 days in the context of small molecules. ~3000 small molecules were used, each at 3 separate concentrations (0.08uM, 0.3uM, 3uM) across twenty-seven 384-well plates. Each plate contained replicate control wells. 2) RRASL-seq protocol using cell lysates (n=110 RRASL probe pairs targeting ~50 mRNAs, 9 batches of 3 plates each). 3) Pooling of RRASL-seq amplicons into a single library. 4) Illumina sequencing, n=2, using aliquots from the pooled library. The upper left control wells didn’t have vehicle, because they were unstimulated t=0. The lower left control wells did have vehicle.

Briefly, RASL-seq assays rely upon RNA-templated ligation of oligonucleotide probe pairs that anneal to complementary sequences^4^. RRASL-seq utilizes T4 RNA Ligase 2 (Rnl2), which requires two ribonucleotide bases in the 3’OH position of the RRASL acceptor probe and a donor phosphate on the 5’ end of the RRASL donor probe. After annealing, excess RRASL probes are washed away, increasing the specificity of the ligation reaction. Each RRASL probe pair is designed to contain flanking adaptor sequences, allowing addition of well and plate-specific barcodes during the PCR step. Samples are then pooled for sequencing. This protocol yields a defined structure for the Illumina RRASL-seq FASTQ reads: well barcode (8 nucleotides), adaptor 1 (17 nucleotides), ligated RRASL probe pair sequence (~40 nucleotides), adaptor 2 (17 nucleotides), and plate barcode (7 nucleotides). This defined structure reduces the complexity of the alignment process and facilitates the analysis of observed error profiles.

For some applications the quantitative accuracy of RRASL-seq probe mapping does not affect the results (e.g. viral RNA detection). However, highly accurate RRASL probe alignment, barcode demultiplexing, and data normalization are essential for any use case that requires transcript abundance estimates to be compared across barcoded samples (e.g. high throughput screening). Additionally, given the cost-efficiency of RASL-seq, computational methods that provide timely analysis of thousands of samples are needed for alignment, demultiplexing, and normalization.

Here, we describe computational methods to design and analyze RRASL-seq experiments that have been implemented as an open source software package, RASLseqTools. Using sample data from a large RRASL-seq experiment, we provide empirical estimates of experimental, sequencing, and alignment error that offer guidance for planning and analyzing RRASL-seq experiments. In addition, we describe visualization strategies and five different normalization methods that minimize systematic error, such as batch effects. Finally, we identify three key predictors of RRASL-seq reproducibility: barcode/probe sequence dissimilarity, sequencing read depth and normalization method.

## MATERIAL AND METHODS

### RRASL-seq Probe Design

One hundred and ten RRASL probe pairs targeting fifty-eight genes were designed using the previously described methodology (Supplementary File 1). Briefly, a probe design protocol similar to Primer-BLAST^5^ was constructed using Python^6^, pandas^7^, Primer3^8^, and MELTING^9^. RRASL probe pairs were designed to complement ~40 nucleotides (nt) of a selected sequence near the three-prime end of each transcript. Each RRASL probe pair was extended 17 nts in the five and three prime directions to accommodate adaptor sequences. Critical probe design parameters included an optimal melting temperature of ~68°C and GC content of ~50%. We define on-target RRASL probe pairs as combinations of donor and acceptor probes designed to measure a target gene. We define mismatched RRASL probe pairs as combinations of donor and acceptor probes not intended to measure a target gene. Additional details and filtering criteria for probe design can be found in the previously published manuscript^3^.

### Human B Cell Experiments

CD19+ B cells isolated from human peripheral blood mononuclear cells were cultured for 7 days in 384-well plates at a concentration of 2,000 cells/well in IgE polarizing conditions (50ng/ml recombinant human IL4 and 20ng/ml megaCD40L), as described previously^3^. Each experimental well contained a single small molecule or natural product at one of three concentrations (0.08uM, 0.3uM, and 3uM). Approximately 3000 small molecules and natural products with known pharmacological activities were selected from the Enzo (n=1730), SelleckChem (n=688), and LOPAC (n=687) chemical libraries (Supplementary File 2).

Each 384-well plate possessed a serial dilution series of control lysates: unstimulated day 0 CD19+ B cell pooled lysates at 250, 500, 1,000, 2,000 cells per well (in duplicate); CD19+ B cells stimulated for 7 days with recombinant IL4 + megaCD40L at 250, 500, 1000, 2000 cells per well (in duplicate), henceforth referred to as Day 0 control lysates and Day 7 control lysates, respectively. Each plate also contained yeast RNA in serial dilution (added to the plates just prior to the RRASL-seq processing), and four empty wells. Finally, each plate contained at least one additional sample of CD19+ B cells stimulated for 7 days with recombinant IL4 + megaCD40L, henceforth referred to as control cells. On day 7, all cells were lysed with Nucleic Acid Purification (NAP; Life Technologies, Catalog no. 4305895) buffer. All NAP lysates were then stored at −80°C until RRASL-seq processing. We refer readers to Larman et al. 2014 for details regarding the RRASL-seq protocol used for these experiments.

### RRASL-seq Sequencing and Basecalling Protocol

The pooled RRASL-seq library was sequenced on two different Illumina HiSeq machines. Sequencing run 1 used 8pM of the RRASL-seq library and 15% phiX library spike-in to balance the base composition. The resulting sequencing run yielded a cluster density of 950,000/mm^2^. Several tiles from this run failed the Illumina quality threshold requiring manual tile selection, a constant phasing matrix, and base calling using the Illumina Offline Basecaller version 1.9.4. Sequencing Run 2 utilized 4.125pM of the RRASL-seq library, 25% phiX library (1.375pM) spike-in to balance the base composition, and 0.5μM of the custom sequencing primer. Casava version 1.8 was used for base-calling for sequencing run 2.

### RRASL-seq Demultiplexing

We designed twenty-seven plate barcodes that have an edit distance no less than two for all pairs of plate barcodes (Supplementary File 2). Plate barcode demultiplexing for sequencing runs 1 and 2 was accomplished using a Levenshtein edit distance (the minimum number of edits required to change one string into another) < 2. Plate barcode error rates were calculated using reads possessing a Levenshtein edit distance of 1 with respect to any single plate barcode. We also designed 384 well barcodes to possess an equal base distribution at each position, in addition to all pairs of well barcodes possessing an edit distance greater than or equal to 3 (Supplementary File 2). Well barcode demultiplexing was accomplished by identifying reads containing a perfect match to a well barcode in the first 8 base calls. We required all reads that did not contain a perfect match to a well barcode be no more than 2 Levenshtein edits, in the first 8 base calls, to a single well barcode. Well barcode error rates were calculated using reads with a Levenshtein edit distance less than three with respect to any single well barcode.

### RRASL-seq Alignment Read Set & Alignment Methods

We randomly selected 200,000 reads from each of the two sequencing runs, henceforth referred to as the alignment read set. We excluded reads with >20% N base calls, more than 2 N base calls in the well barcode region, or missing an exact match to the first 6 base calls of the AD1 adaptor sequence (Supplementary Files 3 & 4). We created a gold-standard alignment call for each FASTQ sequence (Figure 1D) in the alignment read set using exact string matching or Smith-Waterman alignment. We identified FASTQ reads not containing a perfect match to either an on-target or mismatched RRASL probe pair, henceforth referred to as ‘imperfect’ reads and isolated a 45 base-pair substring beginning after the adaptor 1 sequence. Isolated substrings were then aligned to all unique combinations of donor and acceptor RRASL probes (n=11,009) using the Smith-Waterman (SW) algorithm^10^ implemented in the ‘swalign’ python package. The following scoring parameters were used: +2 match, −1 mismatch, −2 gap opening, -2 gap extension. Therefore, we define the maximum alignment score as 2 * probe pair length. We used two criteria to classify ‘imperfect’ read sequences as unmapped. First, ‘imperfect’ reads mapping to multiple on-target or mismatched probe pairs were classified as unmapped. Second, we calculated the Smith-Waterman Difference Score (SW Difference Score), defined as the difference between the maximum SW Alignment Score (maximum SW Alignment Score between read sequence and any combination of RRASL probes) and the SW Threshold (maximum SW alignment score(s) for the mapped RRASL probe pair, excluding self-alignment). Any read with a SW Difference Score less than one was classified as unmapped. We note that the SW Difference Score provides a conservative threshold, yielding a reduction in recall, but an increase in precision. Each mapped read was further classified as aligning to an on-target or mismatched RRASL probe pair.

Reads from the alignment read set were also mapped using the STAR aligner^11^ to a STAR probe database constructed using all unique combinations of donor and acceptor RRASL probes with the following parameters: --runMode genomeGenerate --genomeDir./ --genomeFastaFiles./on_off_probes.fa --genomeChrBinNbits 6 -genomeSAindexNbases 4.

The following STAR alignment parameters were used for read alignment: seedSearchStartLmax 1; seedPerReadNmax 15000; scoreDelOpen −1; scoreDelBase −2; scorelnsOpen −1; scorelnsBase −2; outFilterMultimapNmax 100; outFilterMatchNminOverLread 0.7; scoreGapNoncan −1; outFilterScoreMinOverLread 0.7. Due to the defined structure of RRASL-seq FASTQ reads we specified an integer value to trim five-prime (25 nucleotides) and three-prime base calls (19 nucleotides for Sequencing Run1 reads and 33 nucleotides for Sequencing Run2 reads), in order to remove barcode and adaptor sequences using the clip5pNbases and clip3pNbases command-line arguments. This hard base clipping strategy allows alignment of a small portion of the adaptor 2 (AD2) sequence for any probe pair less than 42 bases long. We filtered all FASTQ read alignments containing sequence identity less than 70%. Multiprocessing of FASTQ read files was accomplished using GNU parallel^12^.

### RRASL-seq Normalization Samples

Reads from the control cells (n=20 per plate), Day 0 control lysates (n=6 per plate), and Day 7 control lysates (n=6 per plate) were used to benchmark normalization methods. Briefly, FASTQ reads were mapped using the STAR aligner and aggregated in the same manner as the alignment read set, creating a matrix of RRASL-seq probe well counts (RRASL-seq probes rows, sample wells - columns). We were most interested in assessing normalization effectiveness in the context of reasonable sampling error, therefore we only considered samples from plates with a median sample library size ≥ 1,000 on-target mapped reads (n=21 plates), as technical precision is poor for samples with a library size lower than this level (Supplementary Figure 2).

### RRASL-seq Normalization Methods

Five normalization methods: Library Size, housekeeping gene (HK gene), median geometric mean ratio (MGMR), median geometric mean ratio by plate (MGMR by plate) and Regularized Logarithm were compared using read count data from the control cells and control lysates. A pseudocount of 1.0 was added to each RRASL-seq probe well count in order to avoid zero division errors. In order to minimize the less interesting effect of low sequencing depth we only considered RRASL-seq probes with a median unnormalized read count > 20 across the control wells and probes targeting immunoglobulin isotype transcripts in the benchmarking experiments (n=64 RRASL-seq on-target probe pairs, Supplementary File 1). However, as described below, the normalization step used all 110 on-target RRASL-seq probe pairs.

Briefly, the Library Size normalized probe count, Y*_i_*„ was calculated by dividing each probe’s read count plus 1 by the corresponding well library size (total on-target mapped reads in the sample plus 1), given by the equation:

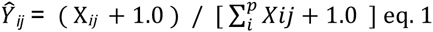

where X*_ij_* is the read count for probe-pair *i* in sample *j*, with *p* indicating all 110 RASLseq probes.

Housekeeping gene normalization was accomplished by dividing each probe’s read count plus 1 by the corresponding well’s read count average for probes NM_001002_RPLP0_909, NM_001101_ACTB_1471, and NM_004068_B2M_313, given by equation:

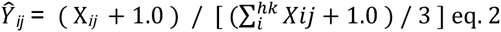

where X*_ij_* is the read count for probe-pair *i* in sample *j*, with *hk* indicating the three aforementioned probes targeting RPLP0, ACTB, and B2M. These probes were selected because they demonstrated the highest pair-wise Pearson’s correlation (across control samples) between probes targeting housekeeping genes.

Median geometric mean ratio, proposed by Anders and Huber^13^, was slightly modified due to the addition of 1 pseudocount to every probe well count in all samples. Briefly, each probe well count was divided by a sample-specific scaling factor, 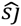, that was determined by taking the median value from the ratio of probe *i* read counts in sample *j* to the geometric mean of probe *i* read counts across all samples under consideration, given by the equation:

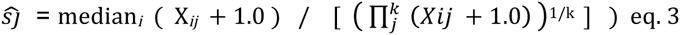

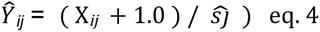

where X*_ij_* is the read count for probe-pair *i* in sample *j*, out of *k* total samples, considering all 110 RASLseq probes. Median geometric mean ratio by plate used the MGMR procedure for each plate separately.

Regularized Logarithm normalization was calculated using the rlogTransformation function in the DESeq2 R package with the Blind parameter set to FALSE, and a design matrix indicating plate membership. Briefly, the rlog function shrinks low read count estimates to the mean probe well count across samples, while returning ~log2 transformed values for high read count probes, we refer readers to Love et al. 2014 for additional technical details^14^.

Normalized RRASL-seq probe well counts were also standardized, defined here as subtracting the average value across samples (mean-centering) and dividing by the standard deviation (scaling). This post-normalization processing step was applied across all samples in the indicated experiment, e.g. standardization of all sixty control cells for the principal component analysis described in Figure 8.

### Visualization of Well Position and Batch Effects

To visualize differences in sequencing depth across plates and reagent batches we plotted the sum of the on-target read counts for each plate grouped by sequencing run, with a color label indicating reagent batch. To visualize well position effects we calculated and plotted the average library size for each well position across plates using the pandas and seaborn python packages. To visualize the variation of sample library sizes stratified by well position we calculated and plotted the coefficient of variation across plates using the pandas and seaborn python packages.

### RRASL-seq Normalization Benchmarking

Once the normalized probe well counts were calculated, we used several approaches to assess normalization effectiveness. Our control cells are technical replicates confounded by reagent batch, plate, and sequencing read depth and therefore useful for normalization benchmarking. We utilized the control cells to assess the effects of five normalization methods using relative log expression (RLE) plots of housekeeping genes, Pearson’s correlation between immunoglobulin RRASL-seq probes and principal component analysis.

For each control cell sample we constructed boxplots of the relative log expression (RLE) of RRASL-seq probes (Supplementary Figure 3). RLE was computed by dividing the RRASL-seq probe well count by the median RRASL-seq probe well count across samples. Samples with a central tendency deviating away from 0 were considered biased by systematic error.

To assess the dependence of normalization on sequencing read depth and global gene expression patterns, we used all control cells to compute the first and second principal components, and then plotted these components against sample library size. Sample library size is defined as the sum of on-target RRASL-seq probe counts in a sample, and is the value used for the Library Size normalization method.

We also computed and plotted the Pearson’s correlation between RRASL-seq probes targeting immunoglobulin transcripts across all control cells. We judged each normalization method using a signal-to-noise score, defined as the difference between the average Pearson’s correlation for probes targeting the same isotype and the average Pearson’s correlation to probes targeting different isotypes (e.g. average pair-wise correlation between the probes targeting IGHA minus the average correlation between probes targeting IGHA and all probes targeting immunoglobulin isotypes other than IGHA).

We used generalized linear models (GLM, negative binomial distribution, log-link function), implemented in the statsmodels python package, to assess the reduction in library size bias. We constructed two nested GLMs for each RRASL-seq probe. The first model predicts RRASL-seq probe count as a function of plate, batch, and sample library size, and the second model predicts RRASL-seq probe count as a function of plate and batch. We counted the number of GLMs that produced a statistically significant coefficient for the library size predictor using unnormalized or normalized data. We then calculated the difference in variance explained between the two models to assess the influence of library size on prediction accuracy.

To further evaluate the usefulness of the five normalization methods we leveraged the 2000 cell, 1000 cell and 500 cell Day 0 and Day 7 control lysates. We restricted our analysis to plates with a median sample library size above 1000 reads (n=21 plates). Furthermore, we only used the technical duplicate with the higher sample library size because we observed a strong relationship between sample library size and probe count correlation for technical duplicates on the same plate (Supplementary Figure 3).

Principal component analysis of Day 0 and Day 7 serial dilution cell lysates from three plates was carried out, as described for the control samples, in order to evaluate whether normalization methods can produce both a clear separation between the biological conditions in the first principal component.

Fold changes (Day 7 / Day 0) for each probe were calculated after normalization, but before log2-transformation, using data from cell lysates on the same plate. Box plots of fold changes (across plates) were plotted for several housekeeping genes. The standard deviation of the log2-transformed fold change across the twenty-one plates was computed for each of the 64 RRASL-seq probes. To assess the relationship between cell density and fold change precision we compared the log2-transformed fold-change vectors and the standard deviation of the log2-transformed fold-change vectors from the 2000 cell and 500 cell conditions using paired t-tests. A paired t-test was used in order to control for variance inherent to each probe.

### RASLseqTools Design Pattern

RASLseqTools has been implemented in Python^6^ with extensive use of the following open source python libraries: SciPy^15^, NumPy^16^, Pandas^7^, swalign, Levenshtein edit distance, seaborn^17^, and Matplotlib^18^, in addition to the STAR^11^ and BLASTn^19^ aligners. RASLseqProbes requires a tab-separated input file describing the donor and acceptor RRASL probe pairs along with the adaptor sequences used for PCR barcoding. RASLseqProbes subsequently creates a probe database specific to the user-selected alignment method (currently BLASTn or STAR). A function is available to calculate the minimum Levenshtein edit distances between all combinations of RRASL probes. Levenshtein edit distance is helpful during barcode and RRASL probe selection prior to experimental execution, as this distance is a major determinant of unambiguous barcode demultiplexing and probe mapping. We have also implemented several visualization and normalization methods as convenience modules in RASLseqTools.

## RESULTS

### RRASL-seq B cell Screen

We recently conducted a RRASL-seq cell-based screen of ~10,000 conditions in 384 well plates, run as 9 separate reagent batches (Figure 1). A final, pooled PCR library was subjected to two separate sequencing runs on two different Illumina HiSeq instruments. Briefly, human CD19+ B cells were stimulated with rhIL4 and megaCD40L +/− compounds from three chemical libraries with pharmacological activity (Enzo, SelleckChem, and Lopac) for 7 days in 384-well plates. Each sample well contained a spike-in quality control polyadenylated RNA sequence (from M13 phage), and was assigned a 7 nt plate-specific barcode and an 8 nt well-specific barcode. In addition, each plate contained Day 0 or Day 7 stimulated B cell control lysates plated in duplicate as two-fold serial dilutions (2,000 cells to 250 cells). At least one well (range 1–32) with stimulated B cells was left untreated on each plate. One hundred and ten RRASL probe pairs were designed to assay fifty-eight distinct genes (Supplementary file 1). A subset of these probe sets were designed to measure similar or shared immunoglobulin constant regions.

### RRASL Barcode and Probe Design

Twenty-seven plate barcodes were used to distinguish individual 384-well plates. Twenty-one and six of these plate barcodes possessed a minimum Levenshtein edit distance of 3 and 2, respectively. We have previously described a set of 384 well barcodes that exhibit balanced base composition (Figure 2A), a minimum pairwise Levenshtein edit distance of 3 (Figure 2B), and a variable number of nearest neighbor barcodes with a Levenshtein edit distance of 3 (Figure 2C).

**Figure 2:**
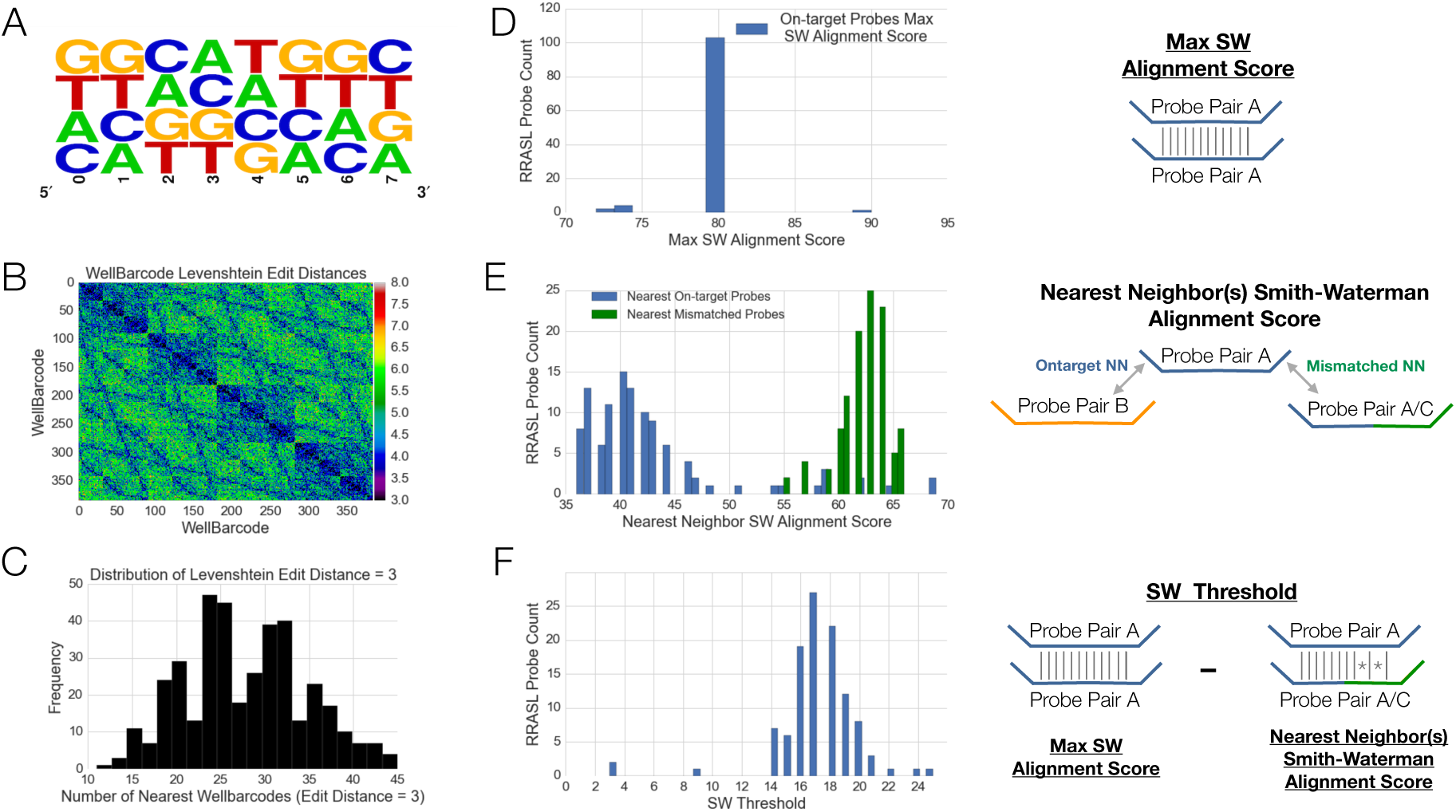
Characteristics of RRASL Well Barcodes and RRASL Probe Pairs: A) Weblogo displaying the position-specific base composition of the 384 well barcodes. B) Pairwise Levenshtein Edit Distances of the 384 well barcodes. C) Distribution of Levenshtein edit distance pairs (edit distance =3) for each well barcode. D) Distribution of maximum Smith-Waterman Alignment Score (Max SW Alignment Score) for 110 RRASL probes used in RRASL-seq screen. E) Distribution of Smith-Waterman alignment scores between on-target RRASL probe pairs and the nearest on-target and off-targetmismatched RRASL probe pair. F) Distribution of Smith-Waterman Threshold scores for 110 RRASL probe pairs.

One hundred and ten RRASL probe pairs were designed to profile fifty-eight distinct genes using our previously published method^3^. The vast majority of the RRASL probe pairs used in this screen had a length of 40 nucleotides, an average melting temperature of 68, and a GC content of 42.4%. We define several useful concepts for Smith-Waterman read alignment classification. First, maximum Smith-Waterman alignment score (max SW Score) is computed as two times the RRASL probe pair length. The vast majority of RRASL probe pairs used in the B cell screen had a max SW Score of 80 (Figure 2D). Nearest Neighbor SW Alignment Score is defined as the Smith-Waterman alignment score for a given probe and its nearest on-target/mismatched probe pair. Figure 2E displays the Nearest Neighbor SW Alignment Score for each of the 110 RRASL-seq probes and their respective nearest on-target (blue) or mismatched (green) probe pair. We find that the majority of nearest sequence neighbors are mismatched RRASL probe pairs, and the remainder target similar or shared immunoglobulin constant regions. Figure 2F displays the Smith-Waterman Threshold Scores (SW Threshold) for each of the 110 RRASL probe pairs, SW Threshold is calculated by subtracting the max SW Score from the Nearest Neighbor SW Alignment Score. The distribution of SW Threshold scores for the 110 RRASL probes is centered at 17 (range 3, 24), indicating clear separation between RRASL probe pairs (Figure 2F).

### RRASL Barcode Demultiplexing

In order to assess the performance of our demultiplexing and alignment strategy, in addition to characterizing the observed error rates, we randomly selected 200,000 reads from each of the two sequencing runs (n=400,000 reads), to create an ‘alignment read set’. We excluded FASTQ reads with excessive ‘N’ base calls or a mismatch base call in the first 8 bases of AD1. All presented percentages assume a denominator of 200,000 reads for the indicated sequencing run.

Summed read counts for the twenty-seven plate barcodes exhibited strong concordance across the alignment read set from sequencing runs 1 and 2 (on-target reads, Figure 3A). We find that 96.72% and 94.3% of the alignment read set possess a perfect sequence match to one of the twenty-seven plate barcodes for sequencing run 1 and 2, respectively. Only 1.54% and 1.18% of the alignment read set reads are more than 1 edit distance away from the expected plate barcodes for sequencing runs 1 and 2, respectively.

**Figure 3:**
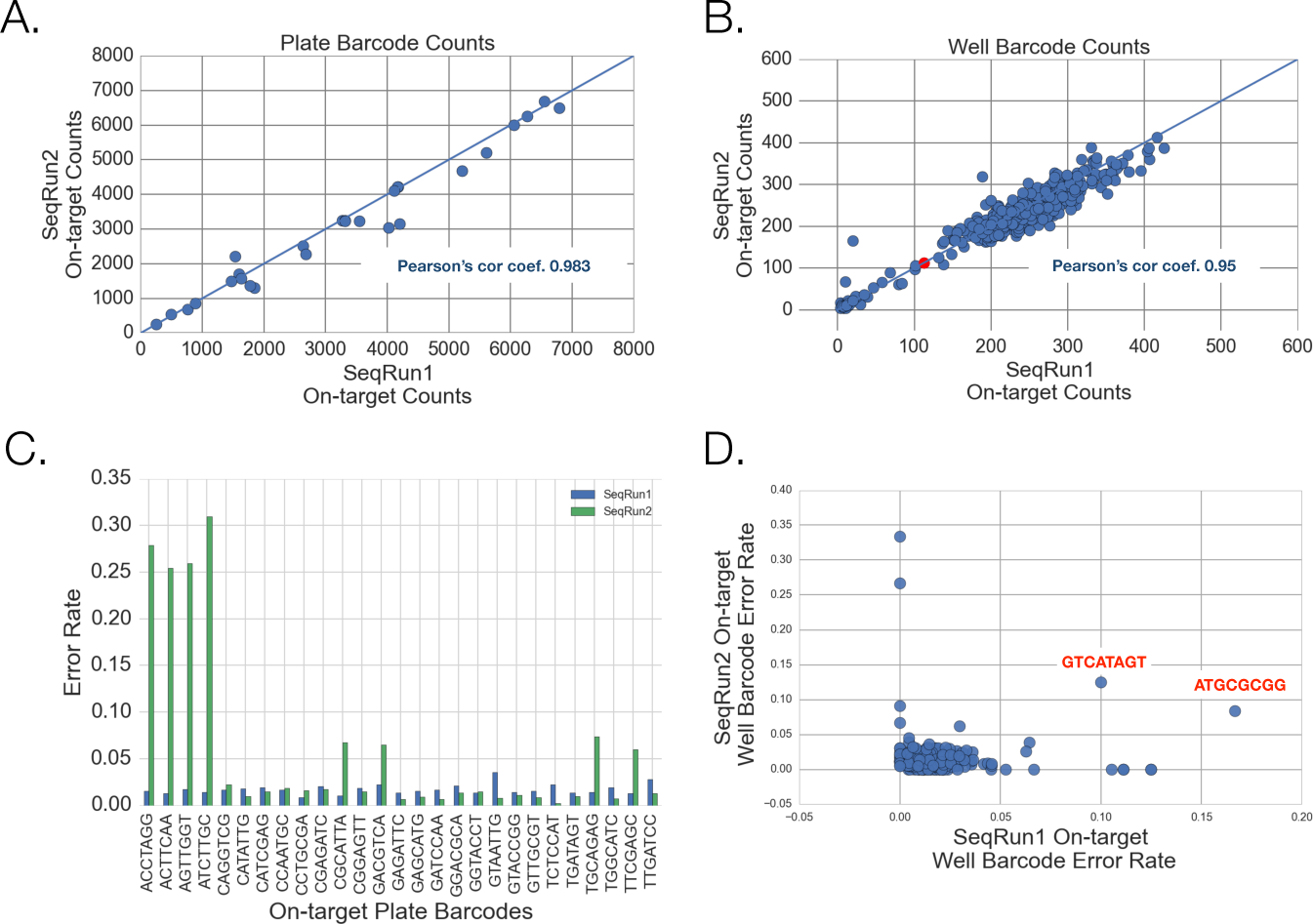
Concordance and Error Rates of Alignment Read Set Barcodes Across Sequencing Runs. A) Scatterplot of on-target alignment read set plate barcode counts across sequencing runs 1 and 2. B) Scatterplot of on-target alignment read set well barcode counts across sequencing runs 1 and 2. AGTAGATC shown in red. C) Error rate of on-target alignment read set plate barcode counts across sequencing runs 1 and 2 D) Scatterplot of alignment read set well barcode error rates in on-target reads from sequencing run 1 and 2. Well barcodes with a consistently higher error rate are labeled.

With respect to well barcode demultiplexing, we observed 97.9% of the first eight bases in the alignment read set mapped perfectly and uniquely to one of the 384 well barcodes for both sequencing runs. 1.65% and 1.48% of the alignment read set well barcodes from sequencing run 1 and 2, respectively, map with one edit distance to a single well barcode. An additional 0.37% and 0. 54% of all RRASL-seq reads possess an edit distance of 2, leaving less than 0.04% of reads unmapped due to errors in the well barcode sequence.

The 384 well barcodes exhibit robust performance in the RRASL-seq assay as assessed by the high Pearson’s correlation, 0.95, in spite of differing base quality between sequencing runs 1 and 2 (Figure 3C). One well barcode, AGTAGATC, has perfect sequence similarity with a standard Illumina adaptor sequence and therefore is ill-suited for RRASL-seq applications if standard demultiplexing methods are used. While most well barcodes exhibit an error rate between 0 and 0.5% (Figure 3D) we have found a subset of well barcodes more prone to either synthesis or sequencing error (Supplementary File 5).

### RRASL-seq Alignment Read Set Probe Mapping

As discussed above, RRASL-seq Illumina libraries are composed of a mixture of the barcoded RRASL-seq ligation product and phiX used to balance base composition. Therefore, our first filtering step identified reads possessing an exact match to the first 8 bases of adaptor sequence, AD1, which excludes PhiX reads. This inclusion criterion removed a percentage of reads corresponding to the phiX fraction spiked into sequencing runs 1 and 2. We found 59.6% and 63.8% of reads in the alignment read set contain a perfect match to either an on-target or mismatched RRASL probe pair (Figure 4A-B), in sequencing runs 1 and 2, respectively. The relatively large and reproducible percentage of mismatched probe alignments indicates the need for further optimization during the RRASL-seq ligation step. A substring (44 bases 3’ to the observed adaptor 1 start position) from each of the remaining ‘imperfect’ alignment read set reads was aligned to all unique combinations of donor and acceptor RRASL probes (n=11,009) using the Smith-Waterman algorithm. Only alignments equal to the highest SW alignment score for each read were retained and used to calculate the SW Difference Score (Supplementary Figure 1B&C). Considering the ‘imperfect’ reads mapping to on-target or mismatched probe pairs, ~23.8% and 20.3%, out of 200,000 reads, mapped to a single RRASL probe pair and ~4.42% and 4.56% mapped to multiple RRASL probe pairs from sequencing runs 1 and 2, respectively. We classified all reads possessing a SW Difference score less than one or mapping to multiple RRASL probe pairs as unmapped, ~16% for both sequencing runs. The average PHRED base quality scores for read positions intersecting and flanking the RRASL probe pair sequence are plotted in figure panels 4C-D for the five read classifications: unmapped, perfect on-target, imperfect on-target, perfect mismatched, and imperfect mismatched. The concordance of the read alignment classification percentages in spite of the large differences in average PHRED scores between the two sequencing runs indicates the Smith-Waterman alignment methods are robust to base calling errors.

**Figure 4:**
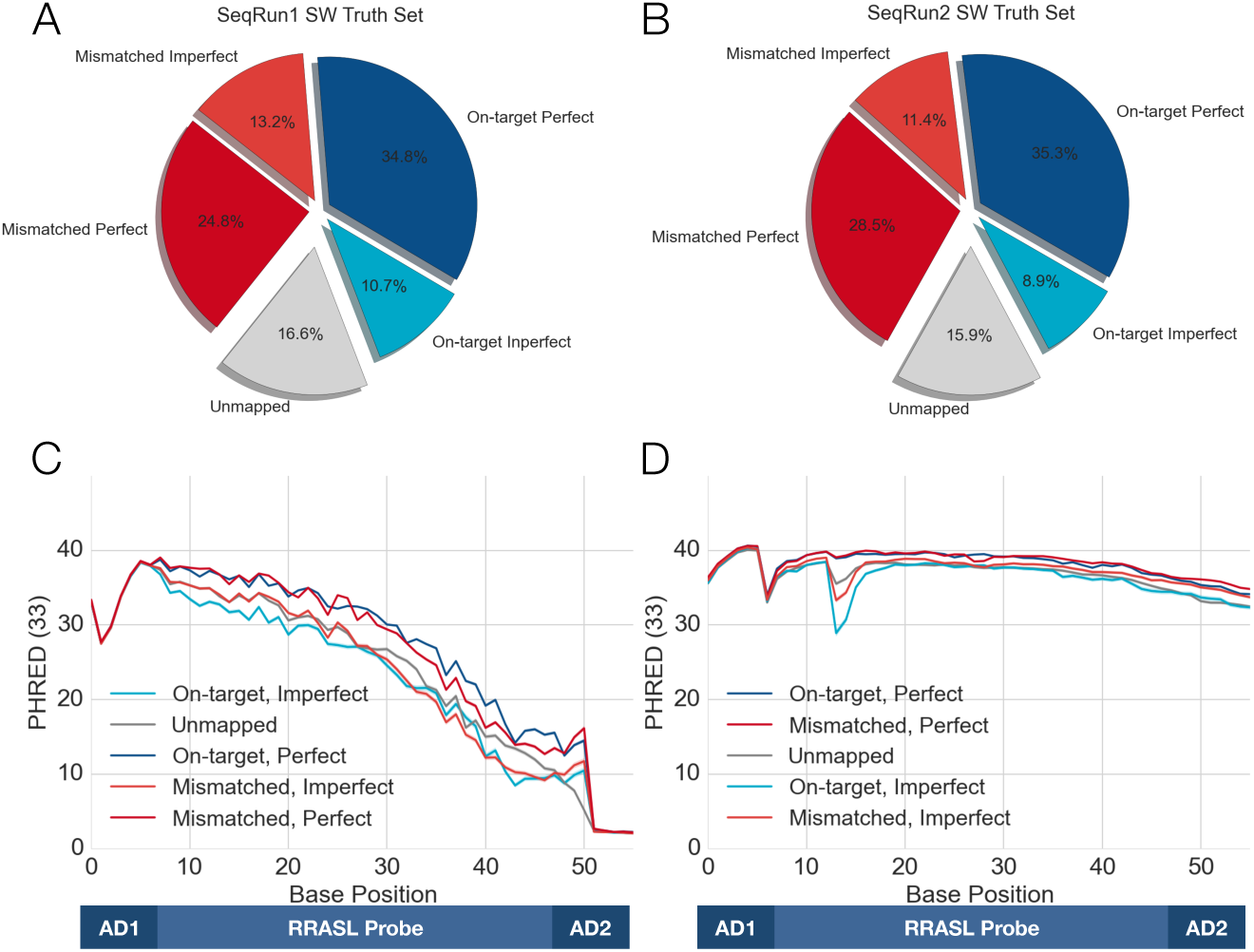
Characterization of Smith-Waterman Alignment Classifications for the alignment read set (n=200,000 reads per sequencing run) across two sequencing runs. A-C) Alignment classifications considered the RRASL probe pair combination (on-target or off-target mismatched), SW Difference Score (<=0 unmapped), and whether the read maps to more than one RRASL probe pair with the maximum SW Alignment Score (>1 unmapped). D-F) Position-specific base quality grouped by alignment classification, average +/− 95%CI. Gray: unmapped, Dark Blue: On-target, Perfect, Light Blue: On-target Imperfect, Dark Red: Off-targetMismatched Perfect, Light Red: Off-targetMismatched, Imperfect.

Figure 5 reports a strong correlation, Pearson’s r ≥ 0.99, between sequencing runs for reads mapping to on-target RRASL-seq probe pairs irrespective of plate or well barcode. Panel C displays the correlation between RRASL-seq probe count estimates for matching and contrasting classification categories. Concordance between reads mapping to mismatched RRASL-seq probe pairs across the two sequencing runs was diminished due to sampling error drop-out, evidenced in the lower left corner of the scatterplot and the observation that the correlation improves when only considering mismatched probes with higher read counts.

**Figure 5:**
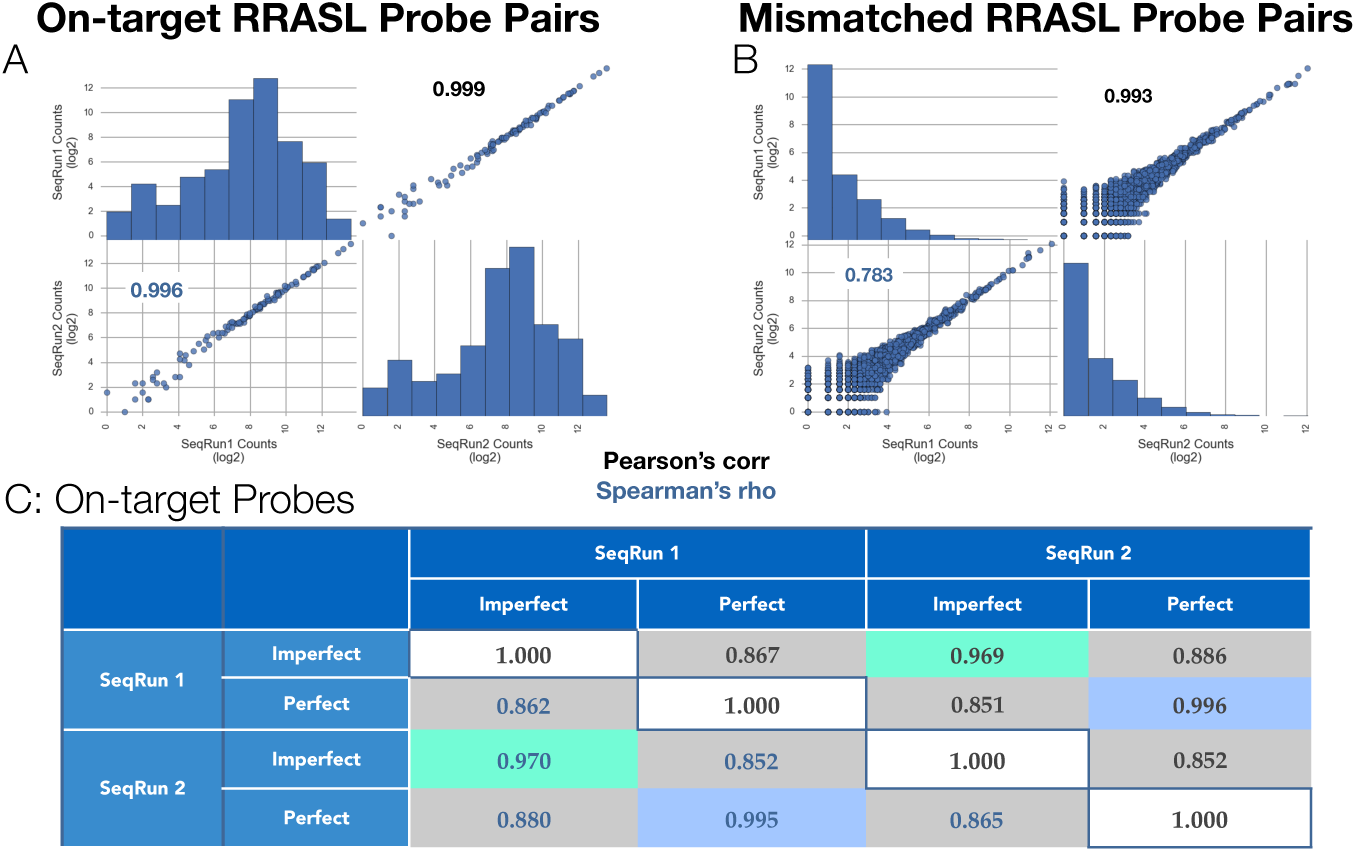
A) Concordance of on-target probe counts in the alignment read set, regardless of plate and well barcode. B) Concordance of off-target mismatched probe counts, in the alignment read set, regardless of plate and well barcode. Table 1: Correlation of perfect and imperfect reads across sequencing runs, irrespective of plate or well barcode. Pearson’s correlation (Black), Spearman’s rho (Blue). Green background color indicates Imperfect-Imperfect comparisons, light blue background indicates Perfect-Perfect comparisons, gray background indicates Imperfect-Perfect and Perfect-Imperfect comparisons.

### RRASLprobe Alignment Benchmarking

The Smith-Waterman algorithm yields optimal pairwise local alignment given an arbitrary scoring scheme, however it is computationally expensive. We benchmarked the STAR aligner using the alignment read set sequences and a reference database composed of all combinations of donor and acceptor RRASL probes (STARrasl). We classified a read as mapped if STAR found greater than 70% sequence identity to a RRASL probe pair. All multi-mapping reads (0.04%, defined as aligning to more than 1 on-target or mismatched probe pair with the same mapping quality score) were classified as unmapped. 43.44% and 43.02% of reads mapped to an on-target RRASL probe pair, and 35.62% and 37.42% of reads mapped to mismatched RRASL probe pairs for sequencing runs 1 and 2, respectively. We observe excellent concordance, 0.999, between STAR and Smith-Waterman alignments (Figure 6 and Table 2), despite the lower number of reads mapped by STAR. We also explored the sensitivity of STAR alignment to flanking bases and found excellent concordance when the expected percent identity filter is adjusted to account for additional flanking bases, i.e. 60% identity when 5 flanking bases are allowed. BLASTn mapping of the alignment read set yields statistically significantly reduced concordance to both Smith-Waterman and STAR alignments (~0.9), and requires at least 300% more wall time. Finally, we used the STAR aligner to map the alignment read set to the Human Genome reference (hg19), which also yielded excellent concordance, 0.999, with both the STARrasl and Smith-Waterman alignment results.

**Figure 6:**
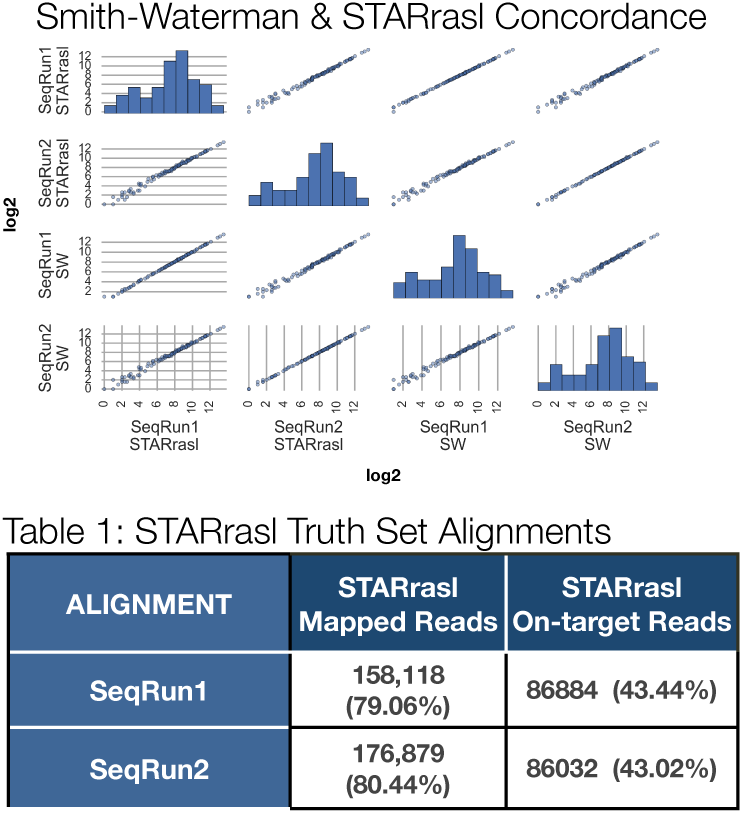
STAR Alignment Statistics (Table 2). Scatterplot comparing the Smith-Waterman and STARrasl algorithms for mapping of the alignment read set. Each point represents a single RRASL-seq probe pair (n=110)

### Visualization of Systematic Bias

Batch and/or well position effects may confound RRASL-seq experiments requiring inter-plate and/or intra-plate comparisons. We have implemented several visualization methods to identify these systematic errors. The first visualization for identifying batch effects revealed dramatic differences between the total on-target mapped read counts for each plate and batch using alignment read set data (Figure 7A). The second visualization displays heatmaps of average sample library size and coefficient of variation (CV) of sample library sizes across plates by well position (Figure 7B&C). An experiment with minimal well position effects will show a uniform average and a low CV across the plate. However, we observe substantial variation in average sample library size (well position effects) and coefficient of variation (batch effects) in the first four rows and edge columns, respectively, indicating systematic bias within plates. The column effect may be due to contamination of columns 1 and 2 during incubation, which prompted the investigators to remove media from these wells prior to lysis. A similar analysis using all reads from the RRASL-seq screen yields the same pattern.

**Figure 7:**
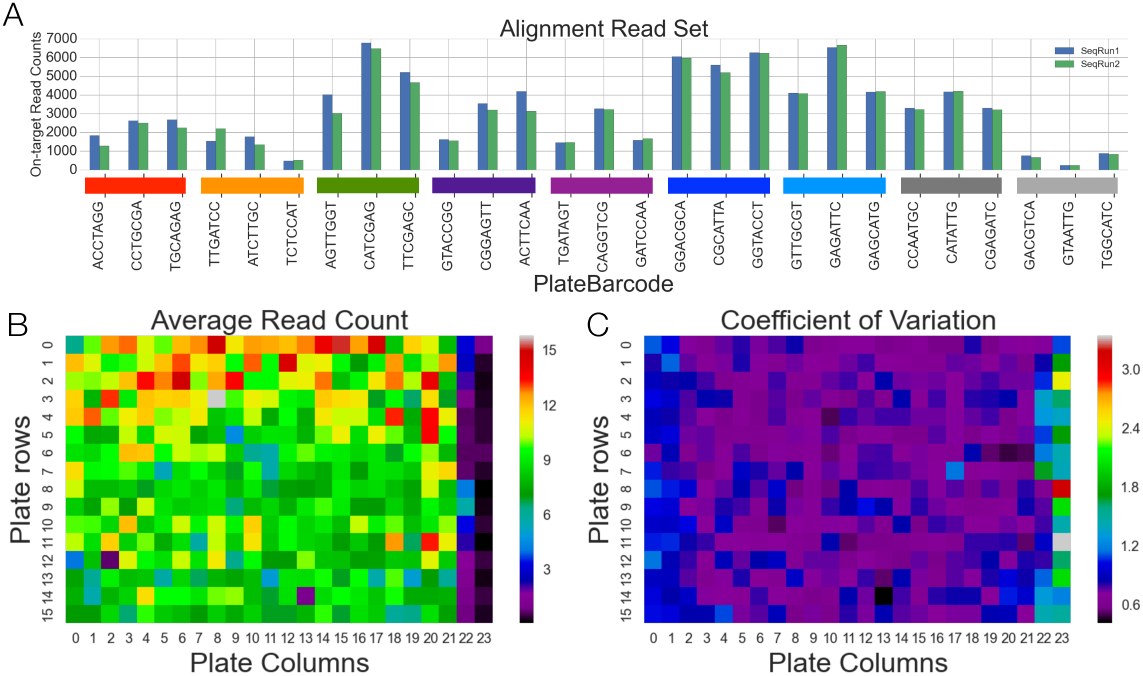
Visualization of Systematic Bias. A) Total on-target read counts by plate, batch, and sequencing run from the alignment read set. Color bars indicate reagent batch. B) Mean well library size across plates. B) Well library size coefficient of variation across plates

### Methods to Correct Systematic Bias

Given the systematic bias detected from the aforementioned visualization methods, we sought to evaluate more advanced visualization and normalization methods using the sixty control cell technical replicates, in addition to using Day 0 and Day 7 samples from the control lysates. We first consider the sixty control cell technical replicates.

Principal component analysis^17^ (PCA) has been used extensively to assess systematic error in gene expression experiments as it captures higher order structure in low dimensional space^18^. We used STAR aligned on-target RRASL probe counts (n=64 RRASL-seq probes) from control cell technical replicates on three separate plates (20 control samples per plate, Figure 8A). Normalized probe counts were mean-centered and scaled to have a standard deviation of 1 across all sixty samples before fitting the principal component model. Technical replicates with minimal systematic bias should display a random array of well positions in 2D PC space and produce principal components explaining little variance. However, we see marked clustering of wells by plate/batch and principal components explaining a large amount of variation, >70% variance explained for the first PC (Figure 8 left panel) when using unnormalized data.

**Figure 8:**
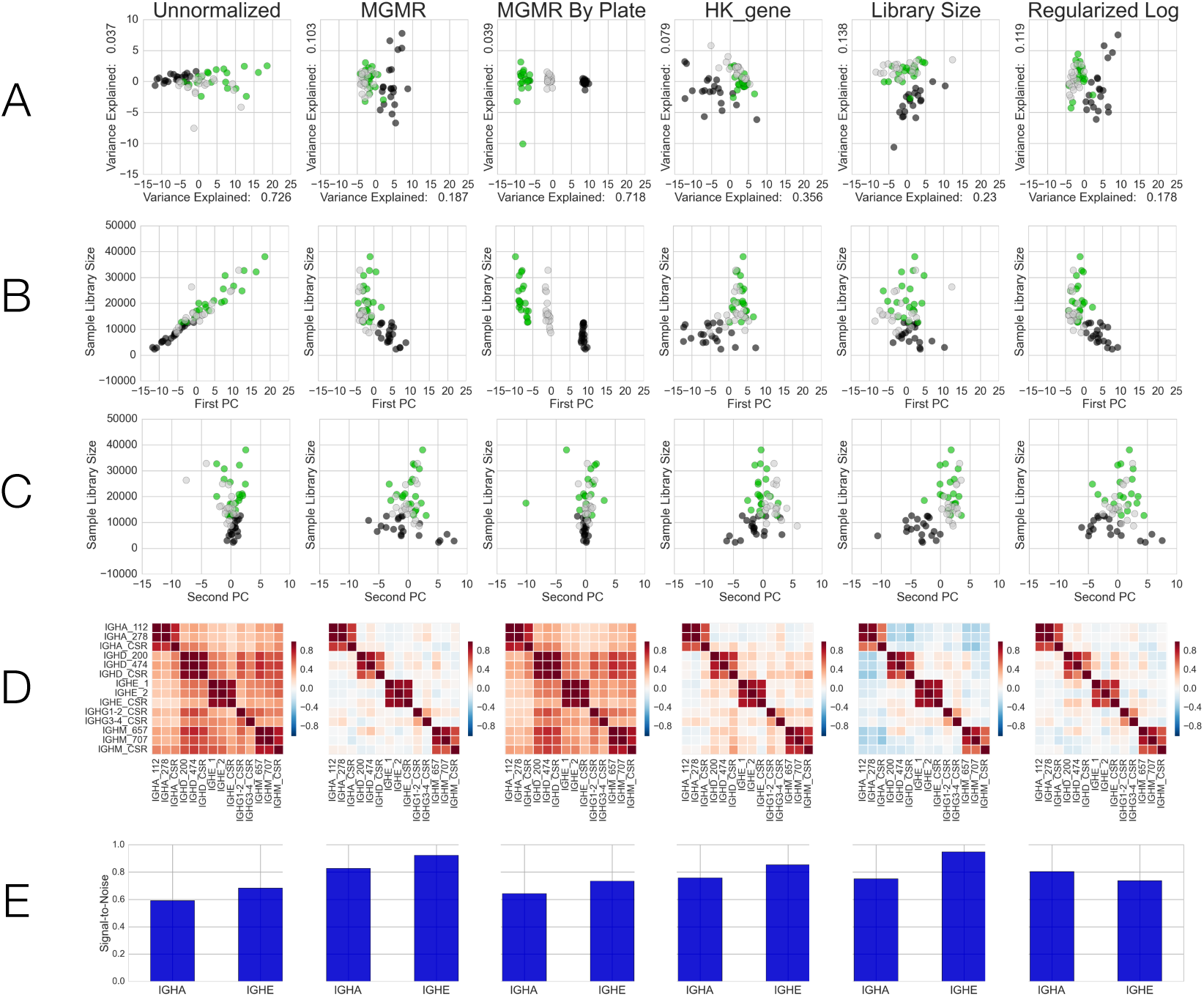
Control Sample Normalization Analysis. A) Scatterplot of the first two principal components using control sample data (n=60 samples, 20 per plate, color indicates plate). B) Scatterplot of the first principal component vs library size for each of the sixty control samples. C) Scatterplot of the second principal component vs library size for each of the sixty control samples. D) Heatmap of pairwise Pearson’s correlation between immunoglobulin isotypes across the sixty control samples. E) Signal-to-noise score for RRASL-seq probes targeting IGHA and IGHE. Signal-to-noise score is defined as the difference between the average Pearson’s correlation for probes targeting the same isotype and the average Pearson’s correlation to probes targeting different isotypes. RASL-seq probe count data was normalized using the indicated method and subsequently mean-centered and scaled to have a standard deviation of 1.

We hypothesized that the large discrepancy in sample library size was a primary confounding factor and would be tightly associated with the first principal component. Principal component regression of the unnormalized probe counts revealed sample library size as the strongest driver of variance both within and across the sixty control cell samples, Pearson’s correlation between first principal component and sample library size was greater than 0.95, Figure 8B. We observe weak correlation between sample library size and the second principal component in the unnormalized PCA model.

Five normalization methods: median geometric mean ratio (MGMR), median geometric mean ratio by plate (MGMR by plate), housekeeping gene (HK gene), Library Size, and Regularized Logarithm were applied to read count data from the control cell samples. All normalization methods, except MGMR by plate greatly reduced the variance explained by the first principal component and minimized the confounding effect of sample library size.

The principal component analysis suggests that the normalization methods are reducing systematic error introduced by library size discrepancies; however an alternative explanation is that the normalization methods are destroying the correlation between features arising from true differences in biological condition. In order to test whether the normalization methods preserve and enhance observed associations between features, we plotted the Pearson’s correlation between probes targeting immunoglobulin isotypes using the sixty control samples (Figure 8D). Visual inspection revealed a dampening of off-diagonal associations. We computed a signal-to-noise score by calculating the difference between the average Pearson’s correlation for probes targeting the same isotype and the average Pearson’s correlation of those probes targeting different isotypes (Figure 8E). We find every normalization method improves the signal-to-noise score, with Library Size and MGMR normalization yielding the largest increase over unnormalized data (Supplementary Table 1) indicating these methods preserve and enhance true associations between probe features. The poor performance of MGMR by plate is likely explained by batch/plate effects driving spurious correlations, a hypothesis supported by the finding that standard scaling each plate individually yields much higher signal-to-noise scores for MGMR by plate.

We assessed the reduction in library size bias by constructing nested generalized linear models (GLM) to predict on-target RRASL-seq probe counts (n=64 RRASL-seq probes) in the control samples using plate, batch, +/− sample library size as predictors. We counted the number of GLM models yielding a statistically significant library size coefficient from the full model when using unnormalized or normalized probe count data. Fifty-eight out of sixty-four GLM models using unnormalized probe counts yielded a statistically significant (Bonferroni-corrected) coefficient for the library size predictor. Only two probes yielded statistically significant coefficients for the models trained using normalized data. We also found that library size explained, on average, 31% of the variance for the corresponding RRASL-seq probe count distribution when using unnormalized data. Library size explained, on average, less than 3% of the variance for the corresponding normalized RRASL-seq probe count distribution (Supplementary File 6).

We further explored the usefulness of the five normalization methods using the control lysates, which are biological and technical replicates confounded by reagent batch, plate, sequencing read depth, and cell density input. The control lysates are composed of unstimulated Day 0 CD19+ B cells or Day 7 CD19+ B cells stimulated with rhIL4 + megaCD40L from the same three plates as the cell control samples. STAR-aligned RRASL-seq well probe counts from the serial dilution series (2000, 1000, 500, cells per well in technical duplicate) were used for the following normalizastion analyses.

Principal component analysis using data from the control lysates was performed in a similar fashion to the cell sample controls (Figure 9A). We found weak discrimination between the biological conditions (Day 0 vs Day 7) along the first principal component using unnormalized data. We expected well performing normalization methods to produce random mixing of serial dilution cell densities within two separate biological condition clusters, in principal component space. We found that every normalization method, except MGMR by plate, enhances the discrimination between Day 0 and Day 7 cell lysate controls, as can be seen in the clear clustering by biological condition on the x-axis. Interestingly, library size bias becomes more prominent in the second principal component of the normalized data, particularly for the Day 0 controls.

**Figure 9:**
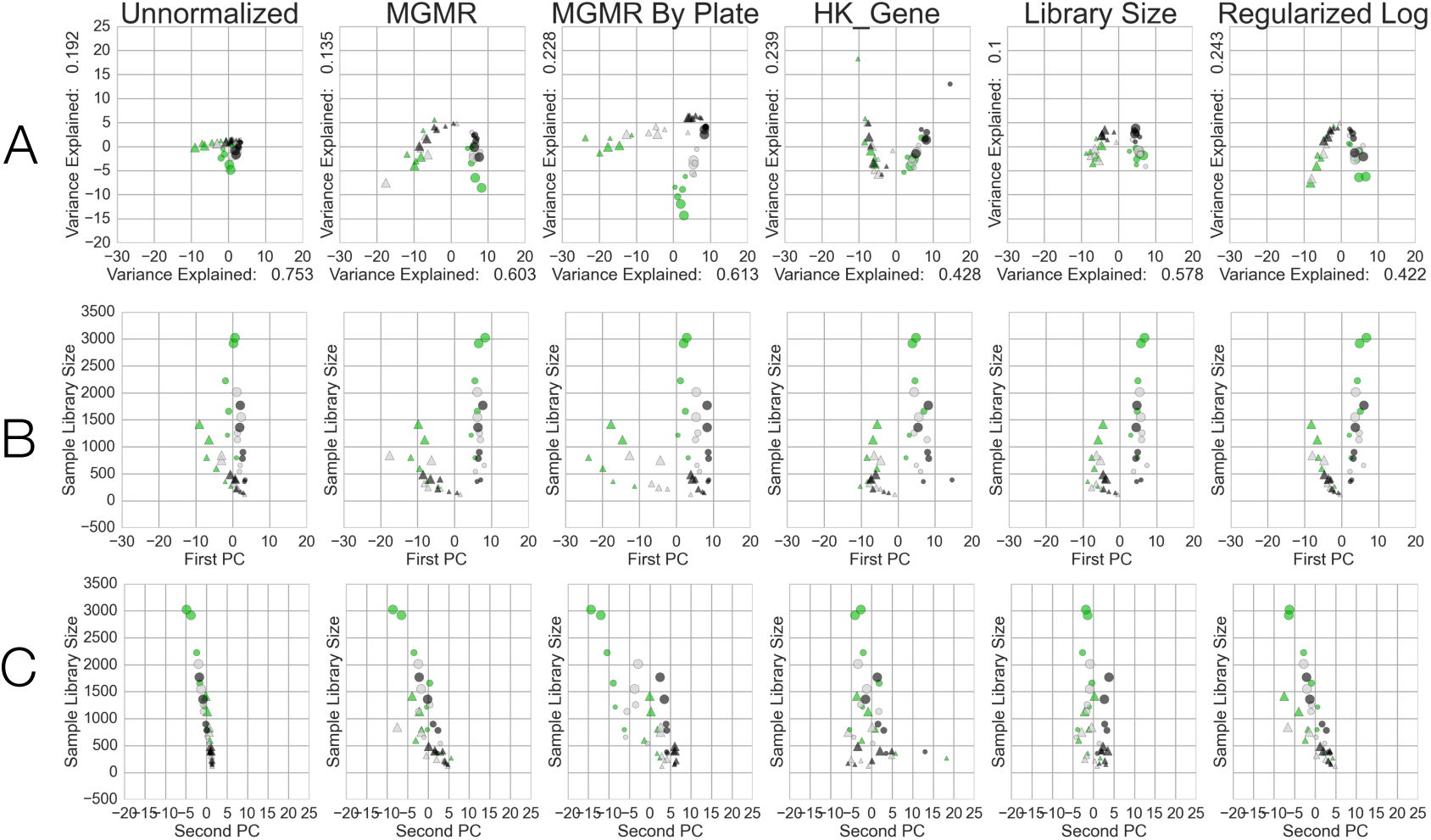
Principal Component Analysis of Normalized Cell Lysate Control Data. A) Scatterplots of the first two principal components of Cell Lysate Controls (x-axis first PC, y-axis second PC). Day 0 Cell Lysate Controls (2000, 1000, 500 cell lysate input samples in duplicate from three separate plates, triangles, n=18) and Day 7 Cell Lysate Controls (2000, 1000, 500 cell lysate input samples in duplicate from three separate plates, circles, n=18) were normalized according to the indicated column header. Each vector of normalized probe counts was then mean-centered and scaled by the standard deviation before PCA. Note: size of marker indicates cell lysate input (largest marker = 2000, smallest marker = 500). Library Size and Regularized Logarithm produce the clearest improvement over unnormalized data for separating Day 0 from Day 7 Cell Lysate controls. B) Scatterplot of first PC value (x-axis) and well library size (y-axis) for each sample. All methods display a correlation between Day 0 library size and the first PC, unsurprisingly, Library Size normalization offers the greatest reduction for the association. C) Scatterplot of second PC value (x-axis) and well library size (y-axis) for each sample. All methods, including no normalization, produce a clear association between well library size and the second PC, indicating well library size remains a significant source of bias after normalization.

One well-established method of systematic bias correction is the use of fold change, calculated by computing a ratio of normalized sample counts to an appropriate control. A well-matched negative control, i.e. Day 0 cell lysates, should produce similar magnitude and low dispersion fold change estimates across serial dilution cell densities and plates/batches. Specifically, we assessed whether the normalization methods corrected for both input RASL-seq library size and sequencing depth in the context of fold change estimation. We hypothesize that effective normalization methods will center the fold-change estimates near zero for housekeeping genes and that the variance in the fold-change estimates will be inversely related to the cell density.

We first investigated the effect of normalization on the central tendency of the estimated log2-transformed fold change (Day 7 / Day 0) for RRASL-seq probes across serial dilution cell densities and plates using data from the control lysates. Boxplots of fold changes for RRASL-seq probes targeting housekeeping genes are shown in Figure 10A. Visual inspection of the boxplots reveals unnormalized data produces log2 fold-change estimates shifted away from zero for several housekeeping genes and also displays tighter variance in the 2000 cell density condition. To formally test the expectation of similar central tendencies between the 2000 and 500 cell density conditions we calculated the number of RRASL-seq probes displaying a statistically significant Bonferroni-corrected paired t-test for unnormalized data and after the application of each normalization method (Supplementary Table 2). Unnormalized data produces an unacceptable 61 statistically significant probes out of 64. All normalization methods yielded dramatically fewer statistically significant probes. Interestingly, although Regularized Log normalization produced the smallest variance, which is unsurprising given the methods shrinkage of low count probes to the average count across samples, it yields 26 statistically significant probes, which is far more than the two, three, and four statistically significant probes found using housekeeping gene, Library Size, and MGMR by plate normalization, respectively. MGMR normalization produced 19 statistically significant probes.

**Figure 10:**
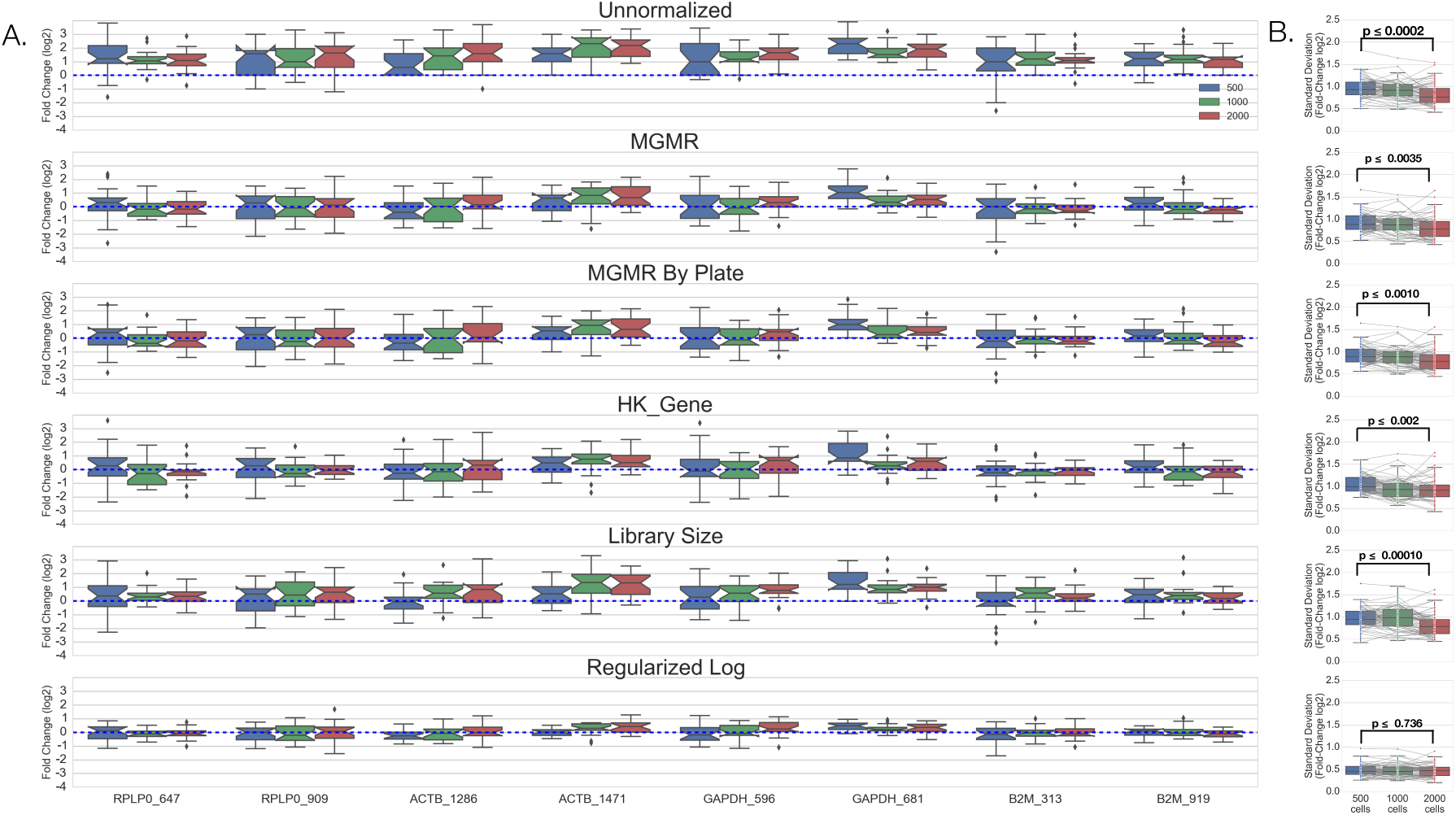
Fold Change Normalization Analysis. A) Boxplots of housekeeping gene log2 fold-changes (Day 7 / Day 0) in control lysate samples grouped by cell density. B) Boxplots displaying the average standard deviation of RRASL-seq probe fold-change across plates (n=64 probes). P-values are shown for a paired t-test comparing the 2000 and 500 cell density conditions

Figure 10B displays boxplots of the observed standard deviations for the sixty-four RRASL-seq probes stratified by cell density and grouped by normalization method. Using a paired t-test, to control variance inherent to different RRASL-seq probes, we find the standard deviation (across plates) of fold changes is statistically significantly lower in the 2000 cell density condition when compared to the 500 cell density condition (Figure 10B). The 2000 cell density samples had, on average, higher read counts, which could bias against the observed finding of lower variance for the 2000 cell density condition given the reported mean-variance association. However, the mean-variance association is likely countered by the increased precision of higher read counts. The exception to this general trend is the fold-change estimates using the Regularized Log normalized data and is likely explained by the replacement of low read count values with the mean across samples. These results offer additional evidence for the primary dependence of technical precision on read depth.

## DISCUSSION

RRASL-seq is a cost-efficient and scalable method for targeted gene expression that leverages multiplexing of both probe sets and barcoded samples. Using data from a high throughput RRASL-seq small molecule screen with a multitude of controls, we explored a variety of factors that may influence the precision of demultiplexing, read alignment, and quantitative comparisons between samples.

We found RRASL-seq offers robust reproducibility despite relatively poor sequencing quality, given sufficient barcode/RRASL probe sequence dissimilarity and read depth. A minimum pairwise Levenshtein edit distance between barcodes is a crucial design parameter, particularly plate barcodes. We recommend an edit distance ≥3 to provide adequate divergence for error correction during the demultiplexing process.

Greater than 80% of RRASL-seq reads containing an Illumina adaptor sequence could be mapped to an on-target or mismatched probe combination, indicating the use of 40 nucleotide RRASL-seq probe pairs with a Smith-Waterman threshold greater than 10 offers ample leeway to correct the relatively low rates of sequencing error common to current Illumina sequencing protocols. In addition, we found the suffix array-based STAR aligner offers dramatically improved computational efficiency and excellent concordance to a conservative Smith-Waterman alignment protocol. We recommend STAR for standard processing of RRASL-seq data.

Any application that requires quantitative comparisons between samples requires identification and minimization of systematic error. We found large differences in sequencing depth across reagent batches, plates, and well positions when visualizing read counts at each of the aforementioned levels.

The large variability of plate-specific read depth is likely a function of pre-sequencing pooling imbalances due to using equal volumes of PCR product, in contrast to equal-molar pooling. Therefore, we recommend assessing library concentration for each plate before pooling in order to achieve a more balanced read depth across plates. Library size imbalances seen at different well positions are likely a function of either robotic pipetting or edge effects during the PCR step. Future experiments will test whether different robotic pipetting strategies change the relationship between well position and library size.

Furthermore, we have shown systematic bias dramatically influences biological inference and must be minimized before quantitative comparisons. We found normalization methods using scaling factors (sample library size or the median geometric mean ratio) offers robust correction for systematic bias introduced by read coverage disparities, well position, and/or reagent batch. More statistically complex transformations such as the regularized logarithm method also reduce systematic error, however we found this particular method very computationally intensive and did not outperform scaling factors in either the signal-to-noise ratio fold change or fold-change central tendency analysis. It is likely that the regularized logarithm method is hampered by the relatively small number of features and may offer superior performance in experiments with far more RRASL-seq probes. While fold-change normalization has a long-standing tradition in experimental biology, we present evidence that normalization prior to fold-change estimation offers dramatic reductions in false discovery rates, even in the context of well-chosen negative controls.

There are several limitations to the presented work. First, although we attempted to design RRASL-seq probes that measure the abundance of a specific transcript isoform, more than half of the probe pairs used in this study bind to multiple isoforms of the target gene. Therefore, we cannot rule out the possibility that alternative splicing has influenced probe counts for RRASL-seq probe pairs that can bind to multiple isoforms. Measuring all known splicing junctions for each target gene is an elegant and useful strategy to overcome this limitation^22^. Second, outliers can produce spuriously large correlation coefficients and can undermine the use of correlation statistics as a measure of reproducibility. The presented scatterplots provide strong evidence that the reported correlations are an accurate reflection of reproducibility in the context of differing library sizes. Third, we recognize that a design matrix can and likely should be employed to further minimize batch effects for any formal statistical comparison between samples, however these investigations are beyond the scope of this manuscript. Finally, we have omitted consideration of robust statistics (e.g. median absolute deviation) in the current work, but acknowledge their utility in the context of large sample sizes enabled by the cost-efficiency of RRASL-seq. The utility of robust statistics will be presented with the comprehensive analysis of the B cell screen in a subsequent manuscript.

To facilitate the use of RRASL-seq we have developed an open source python package, RASLseqTools (https://github.com/erscott/RASLseqTools), offering access to methods useful for the design and analysis of RRASL-seq data. RASLseqTools offers methods to: 1) assess the sequence similarity of barcodes and RRASL probes, 2) demultiplex and align RRASL-seq FASTQ reads, 3) annotate demultiplexed and mapped reads with meta-data, 4) visualize batch effects, and 5) correct systematic error using a variety of normalization methods.

## CONTRIBUTIONS

P.G.S. conceived of and oversaw the project. A.S. oversaw the project. E.R.S. designed and carried out the computational analyses, developed the software, and wrote the manuscript. H.B.L. developed the assays, carried out the experiments, and provided critical feedback. A.T. designed the dual PCR barcodes and helped oversee the project. N.J.S., N.W. and R.T. provided critical assessments of statistical methods. M.W. assisted with software development. R.Z. and L.L.L. designed and carried out validation experiments.

## ACKNOWLEDGEMENT

Chad Locklear (IntegratedDNA Technologies) contributed helpful discussion and the oligonucleotides used in this work. Isha Wallace, Ph.D. (Illumina) contributed helpful assistance with manual tile selection and offline base calling of sequencing run 1 data.

